# Probabilistic mixture models improve calibration of panel-derived tumor mutational burden in the context of both tumor-normal and tumor-only sequencing

**DOI:** 10.1101/2022.04.22.489230

**Authors:** Jordan Anaya, John-William Sidhom, Craig A. Cummings, Alexander S. Baras, AACR Project GENIE Consortium

**Author notes:** Corresponding author: Alexander S. Baras.

## Abstract

**Background:** Tumor mutational burden (TMB) has been investigated as a biomarker for immune checkpoint blockade (ICB) therapy. Increasingly, TMB is being estimated with gene panel-based assays (as opposed to full exome sequencing) and different gene panels cover overlapping but distinct genomic coordinates, making comparisons across panels difficult. Previous studies have suggested that standardization and calibration to exome-derived TMB be done for each panel to ensure comparability. With TMB cutoffs being developed from panel-based assays, there is a need to understand how to properly estimate exomic TMB values from different panel-based assays. Design: Our approach to calibration of panel-derived TMB to exomic TMB proposes the use of probabilistic mixture models that allow for nonlinear relationships along with heteroscedastic error. We examined various inputs including nonsynonymous, synonymous, and hotspot counts along with genetic ancestry. Using the TCGA cohort we generated a tumor-only version of the panel-restricted data by reintroducing private germline variants. Results: We were able to model more accurately the distribution of both tumor-normal and tumor-only data using the proposed probabilistic mixture models as compared to linear regression. Applying a model trained on tumor-normal data to tumor-only input results in biased TMB predictions. Including synonymous mutations resulted in better regression metrics across both data types, but ultimately a model able to dynamically weight the various input mutation types exhibited optimal performance. Including genetic ancestry improved model performance only in the context of tumor-only data, wherein private germline variants are observed.

**Significance:** A probabilistic mixture model better models the nonlinearity and heteroscedasticity of the data as compared to linear regression. Tumor-only panel data is needed to properly calibrate tumor-only panels to exomic TMB. Leveraging the uncertainty of point estimates from these models better informs cohort stratification in terms of TMB.

## INTRODUCTION

Immune checkpoint blockade (ICB) has received FDA approval for several cancer types along with any cancer with mismatch-repair deficiency^1^. However, not all patients respond and some may experience adverse events^2^, necessitating the search for biomarkers that can predict response to ICB^3^. ICB tends to be more effective in cancer types with higher TMB^4^, making it reasonable to explore TMB as a biomarker^5–9^. These studies have found higher TMB to be associated with favorable disease-free survival and assessments of tumor reduction across multiple cancers^10,11^.

Although an appealing metric, TMB values can be influenced by assay type, sample quality, and bioinformatics pipeline^12,13^. Generally speaking, exome sequencing is viewed as the gold standard assay for determining TMB (due to complete coding sequence coverage), but smaller gene-focused assays which cover ≈3% of the exome are often used since they are already part of standard of care to identify potential drug targets. When panel-restriction experiments have been performed with exome data, the panel-derived TMB (panel TMB) values generally correlate well with the exome-derived TMB (exomic TMB)^14,15^, justifying the use of panel data to estimate TMB. Importantly, the use of a panel-based assay for characterizing likelihood of ICB response has been approved by the FDA^16,17^.

Gene panels were not specifically designed to accurately measure TMB; rather, they were repurposed to estimate this measure, and the approval of panel TMB as a biomarker in the context of ICB response has come with some controversy^18^. Panels are biased to include genes commonly mutated in cancer, have higher depth of sequencing than exome data, and often don’t have a paired normal sample from the patient to remove germline variants. Each of these factors biases panels towards reporting a higher value of TMB than would be seen with tumor-normal exome sequencing, using conventional approaches. Furthermore, it is unclear how to properly translate TMB cutoffs determined from a given gene panel assay to another. These issues are widely recognized and an international consortium has been created with the goal of harmonizing TMB values^19^.

In an ideal scenario, each assay would be calibrated using a reference set of samples that contains paired exome and panel sequencing. However, it’s difficult and expensive to generate this data at the scale needed for the variety of assays utilized today. As such, most calibration efforts have been performed by computationally restricting conventional tumor-normal exome sequencing to a given gene panel’s genomic footprint. Using this approach with the tumor-normal exome sequencing from The Cancer Genome Atlas (TCGA), researchers have modeled the uncertainty of TMB estimation that is inherent to subsampling and gene biases^20,21^. But, these efforts have not directly considered differences between germline-subtracted tumor-normal and tumor-only based approaches along with the contributions of differences in read depth between exome and gene panel-based sequencing. Some more recent studies have begun to examine data processing variability by deriving panel TMB estimates from TCGA MC3 variant calls using each laboratory’s specific pipelines^22^.

When TMB is calculated without a matched-normal sample, laboratories rely on various population databases to filter out germline variants. Laboratories may also use algorithms that try to distinguish somatic and germline using the VAFs^23^. Regardless of the approach, there will usually be some error in the ability to accurately identify all germline variants, resulting in so-called “private” germline variants contributing to mutational burden estimates in the context of tumor-only sequencing. This artifact cannot be directly accounted for when using germline-subtracted tumor-normal somatic mutation data such as the commonly used TCGA MC3 data. We suggest the application of models calibrated on germline-subtracted tumor-normal data to tumor-only assays needs to be carefully considered.

Herein we address two of the key sources of bias in gene panel data: (a) regional subsampling of the genome in areas more commonly mutated in cancer, as has been previously characterized (b) artifactual contribution of “private” germline variants to TMB in tumor-only sequencing. Importantly, in contrast to other approaches, we leverage a probabilistic regression framework that can model nonlinearity and heteroscedastic, asymmetric error distributions^24^. At very low TMB panel data initially underestimates exomic TMB, then it overestimates exomic TMB because of the enrichment of genes commonly mutated in cancer, and then at higher values is a closer approximation to exomic TMB. This results in a nonlinear relationship between panel TMB and exomic TMB and complex/non-normal error distributions. Furthermore, we aim to highlight the use of model uncertainty in guiding the interpretation of panel-derived TMB estimates.

## MATERIALS AND METHODS

### Modeling Approach

Our models were constructed in TensorFlow as fully connected neural networks with the general structure of an input predictor vector X followed by 3 hidden layers of size 128, 64, 32 each having softplus activation. The final output layer represents the parameters of the mixture model (means, variances, and mixture weights for each log-normal component), which is then fed into a TensorFlow Probability layer. Each input is represented by a vector ranging from just the panel TMB value to a spectrum of counts across variant types in addition to genetic ancestry. Since the median of the modeled distribution was used as the point estimate, we use mean absolute error (MAE) and Spearman’s rank-order based correlation to measure model performance. Note, models that are monotonically related will report the same rank-order based correlation metrics. The reported model performance metrics are based on the test folds from k-fold cross validation. Additional details of the modeling procedure can be found in the publicly available code repository (https://github.com/OmnesRes/DeepTMB, archived at Zenodo: https://doi.org/10.5281/zenodo.7574237).

### TMB calculation

The TCGA exomic data was generated by different laboratories across multiple years with different technologies and protocols, resulting in variation in the quality of each sample, including depth of sequencing and exome coverage. Given that TMB is defined as mutations per Mb, differences in per sample genomic coverage could be potentially problematic. Instead of assuming a common size of each sample’s captured exome we utilized the available coverage WIGs to give each sample a unique exomic footprint, defining exomic TMB as the number of nonsynonymous mutations per Mb of covered coding sequence. To avoid training with rare, outlier TMB values we limited the regression to include samples up to the 98th percentile of panel-derived input values; as opposed to restricting by the observed exomic TMB values which would artificially truncate the target exomic TMB distributions that we are trying to model.

The TCGA MC3 reports all dinucleotide substitutions as two separate single base substitutions which may individually be reported as silent or nonsynonymous. The Pan-Cancer Analysis of Whole Genomes (PCAWG) consortium previously assumed all of these mutations are in phase^25^; we similarly treated adjacent mutations with similar variant allele counts as being in phase (supplemental methods). We labeled the new merged dinucleotide substitutions as nonsynonymous mutations.

To generate synthetic tumor-only panel data we reintroduced the TCGA PanCanAtlas germline calls into the somatic MC3 calls and applied a spectrum of filters to the combined data with the commonly used gnomAD population databases.

## Data Availability Statement

Publicly available data generated by others were used by the authors. TCGA somatic calls^26^ were obtained from https://gdc.cancer.gov/about-data/publications/mc3-2017. TCGA germline calls^27^ were obtained from https://gdc.cancer.gov/about-data/publications/PanCanAtlas-Germline-AWG. gnomAD annotations^28^ were obtained from https://gnomad.broadinstitute.org/downloads. GENIE 10.1 panel information^29^ was obtained from https://www.synapse.org/#!Synapse:syn25895958. The GFF3 used is available at ftp://ftp.ensembl.org/pub/grch37/current/gff3/homo_sapiens/Homo_sapiens.GRCh37.87.gff3.gz. The Broad coverage WIGs are available at https://www.synapse.org/#!Synapse:syn21785741. The hotspot definitions^30^ were obtained from https://github.com/taylor-lab/hotspots/blob/master/publication_hotspots.vcf. Ancestry information^31^ was obtained from http://52.25.87.215/TCGAA/index.php.

## Software

PyRanges^32^ was used to perform intersections. BCFtools^33^ was used to perform VCF operations. All the analyses are performed in Python 3 and the model was built with TensorFlow 2 and TensorFlow Probability.

## RESULTS

### Model fit

Ordinary linear regression models attempt to model y via a linear combination of X predictors via optimizing the mean squared error (MSE). These linear models make the assumption that the distribution of the residuals are normally distributed and homoscedastic. Figure 1A and 1C show that the assumptions of homoscedasticity and normally distributed error are violated when regressing TMB. Moreover, in Figure 1A and 1C we can appreciate that the “residuals are correlated” in that the linear model predictions tend to slightly overestimate exomic TMB at lower input values and then significantly underestimate at higher input values, violating another assumption in linear regression. These issues are worse in the tumor-only data as compared to the germline-subtracted tumor-normal data (Figure 1C and 1A respectively).

**Figure 1.**
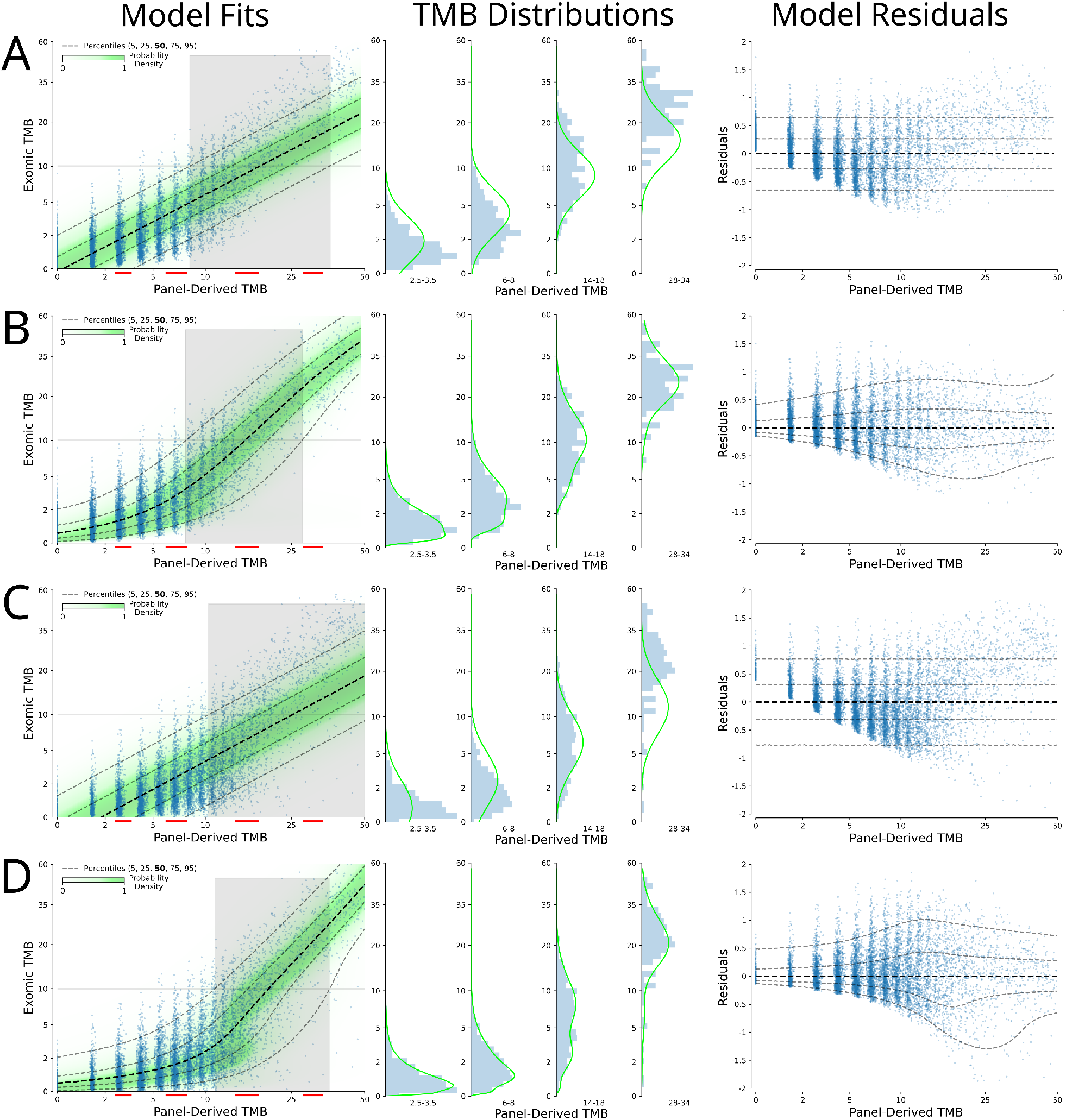
Different modeling strategies for tumor-normal and tumor-only data. Model Fits: Log-log input vs. model output scatter plots with overlaid model distribution probabilities (green) along with the 95% confidence that the model is greater than an exomic TMB of 10 (right of grey area) or less than an exomic TMB of 10 (left of grey area), with red lines indicating what data was used for the histograms; TMB Distributions: Histogram of observed distribution of exomic TMB with overlaid model output distribution at the midpoint of the designated range; Model Residuals: conventional residuals plot. (A) tumor-normal data with linear model, (B) tumor-normal data with proposed mixture model, (C) stringent tumor-only data with linear model, (D) stringent tumor-only data with proposed mixture model.

Minimization of the MSE can be re-posed as minimizing the negative log likelihood of fitting a normal distribution to model the observed y at a given input X, but with fixed variance across all X (as presented in Figure 1A). In our approach we aim to minimize the negative log likelihood of fitting a mixture model of log-normal distributions to estimate the distribution of observed y at a given input X. As can be seen in the second column of Figure 1 (under the “TMB Distributions” header), the observed exomic TMB (y) shows a clear right skew at lower input values and then shifts to a more symmetric, unimodal distribution (albeit not necessarily normal distribution) at higher input values. This highlights the need to leverage more flexible models of regression that don’t depend on the assumptions of ordinary linear regression.

Figure 1A shows the result of fitting a normal distribution (with fixed variance) as a linear function of X to tumor-normal data restricted to the FoundationOne CDx coordinates, which is effectively equivalent to a linear regression that minimizes MSE. The input-output plot shows the data in log-transformed space, allowing for better appreciation of the features of low to intermediate TMB values (exomic TMB < 10). These data would otherwise be compressed and difficult to fully represent and visualize when plotted with rarer high TMB values present in the TCGA data. Linear models of exomic TMB based on panel TMB input poorly estimate the central tendency of the label across the range of inputs and incorrectly estimate the error distribution (as described above).

To better account for the complexities in these data, we model the distribution of the observed exomic TMB at any given input value using a mixture of log-normal distributions. This modeling approach explicitly allows for heteroscedasticity, as can be appreciated in Figure 1B and 1D. This approach better removes any dependency of the residuals on the input values and far better models the distribution of the observed exomic TMB at different input values. The right skew of lower values are well modeled while still allowing for an approximately normally distributed error at higher values. In particular, the effect of private variants on the distribution can be appreciated and characterized in Figure 1D from tumor-only data as compared to germline-subtracted tumor-normal data in Figure 1B.

In considering thresholds based on panel TMB to partition patients into different strata, the common strategy has been to do so based on point estimates from panel-based TMB measures. Such an approach cannot fully account for the uncertainty behind these point estimates of exomic TMB. This will result in particular difficulty in accurately distinguishing values near the chosen threshold. To combat this, one could leverage model uncertainty estimates; but, these need to model the distribution of the data correctly. By better modeling the distribution of exomic TMB at varying input panel TMB, the proposed modeling approach is better suited to leverage model uncertainty than ordinal linear regression.

When considering the exomic TCGA MC3 data and the FoundationOne CDx coordinates, a threshold of 10 for panel TMB results in the following detection rates for true exomic TMB > 10: [a, tumor normal] 53.5% of panel TMB > 10 (n=1486) and 1.1% of panel TMB <= 10 (n=8201), [b, tumor only] 30.6% of panel TMB >10 (n=2758) and 0.5% of panel TMB <= 10 (n=6838). These data show that stratification based on panel TMB with a value of 10 tends to produce a high negative predictive value (NPV) for high exomic TMB (defined by the target of 10 in this context) but poor positive predictive value (PPV). Applying either a linear model or the proposed mixture model and using the point estimates from these models to stratify the data results in improved and more balanced NPV and PPV (NPVs > 95%, PPVs > 80%, Supplemental Table 1).

The use of panel TMB or point estimates from models does not leverage the uncertainty/variability of the data to make a better stratification. Since we have modeled the distribution of y given X, we can stratify the data based on the 95% probability of model output being above or below a given threshold (as opposed to the median or mean of the distribution being greater than a threshold). This is represented in Figure 1 in which the the non-shaded areas show where model output can be said to be greater than a target of 10 with 95% confidence (to the right of the shaded areas) or less than a target of 10 with 95% confidence (to the left of the shaded areas). This results in a tripartite stratification in which the two strata with 95% confidence (roughly 80% of the data) result in NPV/PPV ranging from 95-99% (Supplemental Table 1). Of particular note, the linear model is not able to properly model the distribution of the data and as such its model uncertainty estimates cannot be used to accurately gauge the confidence of the model, and this was particularly evident in tumor-only data. The overall theme of these findings are not specific to the chosen target value of 10.

### Effect of panel inputs

A common approach to estimating nonsynonymous exomic TMB from a gene panel-based assay is to count the number of nonsynonymous mutations detected by the panel and normalize these by the panel size (panel TMB) to arrive at a similar metric as in the exomic TMB (specifically, nonsynonymous mutations per megabase pair). Interestingly, in other approaches for estimating exomic TMB from panel-based assays synonymous mutations may be included and so-called “hotspot” mutations may be excluded. Using our mixture modeling approach (as presented in Figure 1) we investigated how different possible inputs (synonymous/nonsynonymous/hotspots) across 3 different datasets affect model performance in estimating exomic TMB. While not currently employed by any protocols to our knowledge, given that different ancestries have varying degrees of intrinsic germline diversity and representation in the databases used to filter out germline variants from tumor-only data, we also investigated the use of genetic ancestry as an additional input into the model.

Table 1 reports metrics of model performance (MAE and Spearman’s rho) from 5-fold cross validation (for a figure representation see Supplemental Figure 1). As has been well-described previously, panel TMB tends to overestimate exomic TMB (which can be appreciated in Figure 1A, in particular for exomic TMB < 20). In Table 1 we show that applying the proposed mixture model to estimate exomic TMB using nonsynonymous mutations results in a significant improvement in MAE as compared to a linear model. When using the proposed mixture model, removing hotspots improved the MAE of tumor-normal data but interestingly appeared to worsen the MAE of tumor-only data. In contrast, including synonymous mutations universally improved MAE metrics. Including ancestry into the model improved the metrics of tumor-only data but had little effect on tumor-normal data, which is somewhat expected in the context of bona fide somatic mutations of tumor-normal data. Finally, we provided the model with a vector input of counts of all the variant types (nonsynonymous/all mutations/hotspots) along with ancestry allowing the model to contextualize the relative importance of these inputs. This formulation uniformly led to the best regression metrics in cross-validated assessments of performance (Table 1).

**Table 1.** TMB regression metrics for TCGA MC3 samples. Regression metrics for different inputs and models. All metrics are from cross validation. MAE: mean absolute error. MAE and Spearman rank-order correlation (rho) are calculated for samples with panel TMB greater than or equal to 5.

| Data Source | Model | Non-Syn | Syn | Hotspots | Ancestry | MAE | Spearman's rho |
| --- | --- | --- | --- | --- | --- | --- | --- |
| Tumor Normal<br>(matched germline subtracted) | panel TMB | + |  | Included |  | 4.71 | 0.72 |
|  | LM | + |  | Included |  | 3.41 | 0.72 |
|  | MM | + |  | Included |  | 2.83 | 0.72 |
|  | MM | + |  | Removed |  | 2.71 | 0.74 |
|  | MM | + | + | Included |  | 2.45 | 0.78 |
|  | MM | + | + | Removed |  | 2.36 | 0.80 |
|  | MM | + | + | Removed | + | 2.37 | 0.80 |
|  | MM | + | + | Included | + | 2.35 | 0.81 |
| Tumor Only<br>(stringent germline filtering) | panel TMB | + |  | Included |  | 6.60 | 0.62 |
|  | LM | + |  | Included |  | 2.89 | 0.62 |
|  | MM | + |  | Included |  | 2.35 | 0.62 |
|  | MM | + |  | Removed |  | 2.41 | 0.60 |
|  | MM | + | + | Included |  | 2.21 | 0.64 |
|  | MM | + | + | Removed |  | 2.21 | 0.63 |
|  | MM | + | + | Removed | + | 2.16 | 0.64 |
|  | MM | + | + | Included | + | 2.14 | 0.65 |
| Tumor Only<br>(permissive germline filtering) | panel TMB | + |  | Included |  | 8.98 | 0.56 |
|  | LM | + |  | Included |  | 2.77 | 0.56 |
|  | MM | + |  | Included |  | 2.31 | 0.56 |
|  | MM | + |  | Removed |  | 2.36 | 0.53 |
|  | MM | + | + | Included |  | 2.29 | 0.54 |
|  | MM | + | + | Removed |  | 2.32 | 0.52 |
|  | MM | + | + | Removed | + | 2.22 | 0.55 |
|  | MM | + | + | Included | + | 2.17 | 0.58 |

### Effect of tumor-only sequencing

Tumor-only sequencing lacks a matched-germline control, resulting in a variable number of rare/private germline variants that cannot be identified as germline using even the largest population databases or cutting-edge bioinformatic techniques. The inclusion of these private germline variants results in bias (increased number of variants) and noise (variability in the number of increased variants per sample) being introduced in the estimation of the true somatic exomic TMB. Of particular concern given much of the recent literature, if a model is trained on tumor-normal data (such as TCGA MC3) and then applied to tumor-only panel data (which would include private germline variants) then biases of tumor-only data will not have been accounted for, thereby affecting model performance (Figure 2, panel A vs. B and C). To better account for this we constructed a synthetic tumor-only dataset from TCGA data by reintroducing germline variants into the TCGA somatic mutation profiles that could not be filtered out by common strategies across a spectrum of stringencies using the gnomAD 2.1 database, mimicking common filtering processes used with real-world tumor-only data.

**Figure 2.**
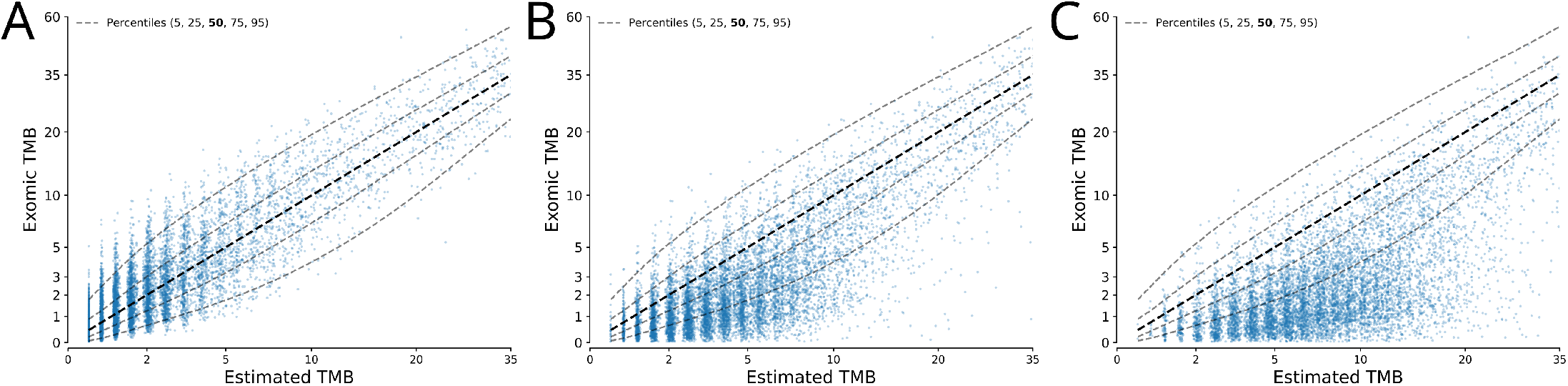
Effect of tumor-only sequencing. Prediction versus true plots for a model trained on tumor-normal data and applied to (A) tumor-normal data, (B) tumor-only stringently filtered data, and (C) tumor-only permissively filtered data.

We used gnomAD 2.1 non-cancer exome and gnomAD 2.1 genome as population databases, and considered the TCGA itself to be a population database of sorts. Using a self-cohort as a population database has the additional benefit of removing potential high frequency artifacts in the data, as is commonly done in the context of individual laboratories. We applied both permissive and stringent criteria for filtering out germline variants using the maximum population specific allele frequency (popmax AF greater than 1% vs. 0.1%) in the gnomAD 2.1 non-cancer exome and gnomAD 2.1 genome population databases in conjunction with the overall allele frequency (AF greater than 0.1% vs. 0.01%). These same popmax AF thresholds were also applied to the TCGA “self-cohort” with known hotspots whitelisted^30^.

Defining somatic mutations as the positive class, our filtering cutoffs resulted in sensitivities/specificities of 99.9%/98.2% vs. 96.5%/99.1% (permissive vs. stringent). For samples used in our regression models we started with an average of 2,451 germline variants per sample in the 4Mb region represented by the union of all panels in AACR GENIE, and after filtering ended with on average 43 and 20 private germline variants per sample for the two filtering stringencies. Studies examining gnomAD have estimated that a given genome has approximately 200 coding variants present at an AF less than 0.1% (private germline variants)^34^. Assuming a 32 Mb coding region, this would equate to roughly 25 variants in a 4 Mb region, which is consistent with our results.

When using the tumor-only dataset we generated to estimate the true somatic, exomic TMB from panel TMB (Figure 1C and D), the introduction of private germline variants results in a bias (artificially inflating panel TMB estimates) and adding noise (increasing the variability of the exomic TMB distribution for a given tumor-only panel TMB value). The proposed mixture modeling approach is able to better fit the central tendency of the data (as can be appreciated when comparing the residuals in Figure 1D vs. 1C); additionally, it is able to better characterize the distribution of exomic TMB, which displays a wider and more asymmetric distribution as compared to tumor-normal matched germline-subtracted data (Figure 1D vs. 1B, respectively).

The number of private germline variants per sample at a given filtering strategy in these data is also dependent on genetic ancestry due to the genetic diversity of different ancestral populations in conjunction with the degree to which they are represented in the databases that are queried (Supplemental Figure 2). As such, we observed slightly different regression models for the different genetic ancestries present in these data (Supplemental Figure 2) and that inclusion of genetic ancestry as an input into the model improved model performance more consistently in the context of tumor-only data (which contain private germline variants, Supplemental Table 2).

### Proposed clinical utility

When deciding whether to prescribe a treatment clinicians must weigh the likelihood of a patient responding vs. the potential risk of the treatment. We suggest that another consideration should be taken into account: the measurement error in the prognostic indicator. To demonstrate what this might look like we leveraged the associated clinical data for the TCGA samples^35^. In this dataset exomic TMB has been shown to be a positive prognostic indicator for bladder urothelial carcinoma (BLCA)^36^, and using these samples we identified the optimal exomic TMB cutoff as 6.7. We used our stringently filtered tumor-only data to train models with the BLCA samples excluded, and as can be seen in Figure 3A although our mixture model was not trained on any of the BLCA samples, its central tendency is a good estimate for the BLCA data and the data is contained within the confidence intervals. In contrast, the linear model’s central tendency is clearly biased and its confidence intervals aren’t applicable (Figure 3B), mirroring what we saw in Figure 1. The inability of the linear model to properly represent the data results in a higher degree of uncertainty (grey area) for being able to confidently (95%) predict a sample as being greater or less than the exomic TMB-derived threshold. In contrast, our proposed mixture model both better estimates the central tendency of this regression and importantly also models the variable degree of uncertainty across the data. This affords our approach a smaller region of data in which it cannot confidently distinguish a sample from the exomic-derived threshold. Ultimately, our model is better able to stratify the observed outcome data based on where each model is 95% confident that the exomic value is greater than the exomic-derived threshold (Figure 3C vs. Figure 3D).

**Figure 3.**
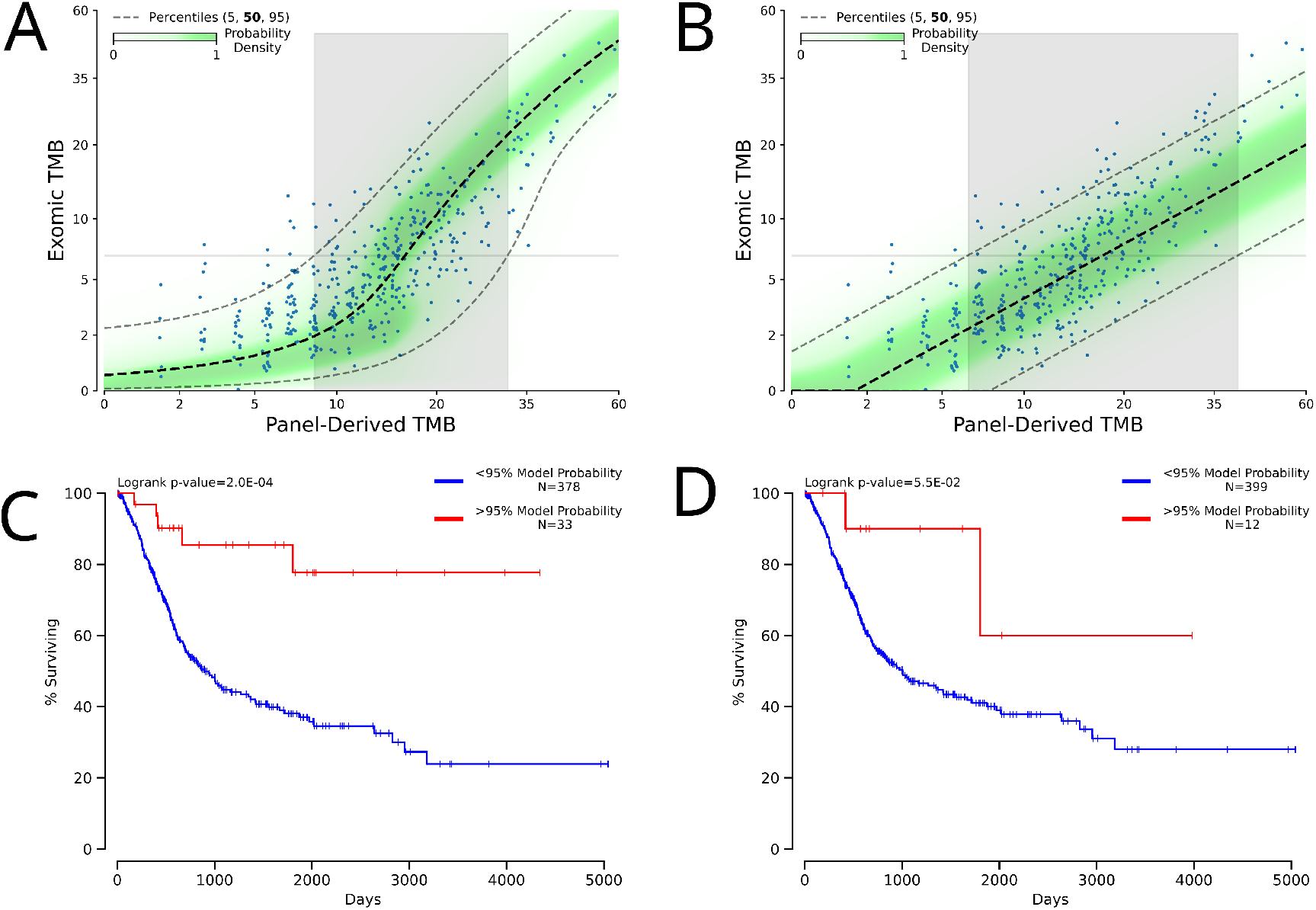
Potential clinical utility of mixture model-derived estimate of uncertainty. (A) Fit of a mixture model with unseen BLCA samples plotted. (B) Fit of a linear model with unseen BLCA samples plotted. In both (A) and (B) to the right of the grey area the model is at least 95% confident the data is greater than the cutoff value of 6.7. (C) Kaplan-Meier survival curves for BLCA samples which the mixture model could say with at least 95% confidence were above the optimal cutoff vs. not. (D) Kaplan-Meier survival curves for BLCA samples which the linear model could say with at least 95% confidence were above the optimal cutoff vs. not.

## DISCUSSION

Given the nonlinear relationship between panel-derived TMB measures and exomic TMB along with the complex error distributions of exomic TMB at different panel-derived TMB ranges, a more flexible modeling approach than linear regression is required to correctly model the data, which we show in Figure 1. In this study we developed a probabilistic model that outputs a probability distribution of the model output (y) for any given input X. From these distributions users can easily get the mode, mean, median or any desired quantile of the probability distribution. Although use of the appropriate modeling strategy is important, it is even more important that every effort be made to ensure the training data resembles the data the model will be applied to. In this study we underscored how significant this can be in the case of tumor-only data and characterized this by reintroducing private germline variants to the TCGA MC3 samples to model tumor-only data.

Previous studies have investigated the performance of different filtering approaches for tumor-only sequencing data, often with the goal of accurately filtering out germline variants^37–44^. Our goal was not to find the optimal filtering algorithm, but to simply use a reasonable spectrum of filtering approaches and characterize the effect that introducing private germline variants would have in the context of calibrating panel TMB measures to exomic TMB. Using the tumor-only data we constructed from TCGA data, the findings would suggest a more stringent filtering strategy would favor more accurate exomic TMB calibration even if that resulted in filtering out some true somatic variants. This is compatible with the notion of increasing the signal (somatic mutations) to noise (germline mutations) ratio given that the target is somatic exomic TMB and there is a large excess of germline variants in relation to somatic ones. One should note however that these somatic variants filtered out under stringent criteria for the purpose of TMB calibration could still be analyzed for other purposes and interpretations, as appropriate.

When using tumor-normal data for calibration there is an inherent assumption that no private germline variants will be present in samples the model will be applied to. However, the majority of academic and commercial labs that perform panel-based somatic mutation profiling use tumor-only approaches, which will result in the inclusion of varying degrees of private germline variants. Population databases alone will unlikely be able to perfectly identify all germline variants from tumor-only data; as such, additional strategies would need to be applied to remove these private germline variants in a manner that does not remove true somatic variants. The tumor-only data we generated from the TCGA using the genomic coordinates of every panel in AACR GENIE constitutes a labeled training data set at the level of the variants (somatic vs. germline) to support the development of such algorithms, which will be the focus of a separate study; the aggregate variant counts used in the current study are available (Data_S1). Interestingly, we show that the amount of private germline variants is dependent on genetic ancestry which has implications on how to estimate exomic TMB from panel TMB.

In addition to often being tumor only, panel data is often sequenced deeper than exomic data. The deeper sequencing has the effect that more mutations will be called with panel data^45^. Additional factors that can affect the number of called mutations include sample purity and clonality^46,47^. While these additional factors would ideally be taken into account when calibrating TMB to a unified standard, the focus of this manuscript was to model the distributions of tumor-normal and tumor-only data, the shape of which will not change if additional mutations are added to both the input predictors and the target.

It is important to not only accurately estimate TMB from panel data, but to also be able to model the uncertainty of such estimates and then use them to inform the use of the model estimates, particularly to consider flagging estimates with high uncertainty. We acknowledge that current thresholds on panel TMB are derived from association to outcome/treatment response and not necessarily correlation with exomic TMB. However, it should be noted that panel TMB around 10 tends to show the highest degree of variability in terms of what the true exomic TMB is. As such, panel TMB thresholds may need to be considered in light of the variability behind them if conceptually they are being considered an estimate of the true exomic TMB, which can be observed exactly if needed. The importance of characterizing uncertainty/variability of estimates underscores the need to use models more expressive than linear models that can account for the nonlinearity and heteroscedasticity present in these data. Our proposed mixture modeling approach serves this purpose well and can allow for a dynamic set of inputs spanning various variant tabulations (synonymous/nonsynonymous/hotspot counts) and sample level properties such as genetic ancestry information as well.

## ACKNOWLEDGMENTS

The results here are in whole or part based upon data generated by the TCGA Research Network.

