## Supplemental Information for "Probabilistic mixture models improve calibration of panel-derived tumor mutational burden in the context of both tumor-normal and tumor-only sequencing"

### TCGA somatic and germline variant processing

Variants in the MC3 public MAF which had a "FILTER" value of either "PASS" , "wga", or "native_wga_mix" were retained. Coverage WIGs were converted to BEDs with wig2bed. Variants were dropped if they didn't fall in the corresponding tumor/normal coverage coordinates. We considered nonsynonymous mutations to be any mutation with the labels "Missense Mutation", "Nonsense Mutation", "Frame Shift Del", "Frame Shift Ins", "In Frame Del", "In Frame Ins", "Nonstop Mutation".

Consecutive single base substitutions were merged into a single mutation if the maximum difference between the average alternative or reference counts and any single alternative or reference count was less than 5 or the maximum percent difference was less than 5%, or the maximum VAF deviation was less than 5%.

### Germline filtering

We used gnomAD 2.1 as the population database for germline filtering. Specifically we used the non_cancer subset of the exome data and all of the genome data. We considered somatic mutations to be the positive label, and as a result defined sensitivity as the number of correctly labeled somatic mutations over all somatic mutations. Specificity was then defined as the number of correctly labeled germline variants over all germline variants.

To construct the synthetic tumor-only dataset the annotation and filtering was applied to both the MC3 calls and PanCanAtlas germline calls and then all remaining calls were treated as somatic. Because germline mutations are always reported as single base substitutions, after annotation neighboring germline mutations were merged as we would merge somatic mutations, with the additional possibility of neighboring germline and somatic SNPs being merged.

### Effect of genetic ancestry

The ability to identify germline variants in the context of tumor-only sequencing is not only a function of the filtering criteria but also determined by the underlying genetic diversity of different ancestral populations in conjunction with the degree to which they are represented in the databases that are queried. In Supplemental Figure 2A, the number of private variants per sample is shown to differ across the TCGA cohort when stratified by genetic ancestry[1]. The findings are similar to what was reported for the gnomAD cohort[2] in that we observed increased private variants in African and Asian ancestries as compared to European. We fit models to each cohort independently in Supplemental Figure 2B and 2C and show therein the model predictions for the different ancestries with stringent and permissive filtering, respectively. In these figures one can observe that the regression lines for non-European ancestries are shifted to the right relative to the European fit; this is due to the additional private germline variants, with the magnitude of the shift roughly correlated to the number of private variants. Indeed, model performance is improved when ancestry is included into the model as characterized in Table 1, but the effect is limited due to most samples being of European ancestry in the TCGA cohort.

To better characterize the effect of including ancestry into the model, we instead performed weighted training to mimic what would happen if each ancestry were equally represented in the TCGA dataset, and looked at the regression metrics per ancestry (Supplemental Table 2). Each ancestry has a different distribution of panel TMB and exomic TMB in both tumor-normal and tumor-only data (Supplemental Figure 3A) and as a result would be expected to display different regression metrics even in the tumor-normal context. Consistent with the observation in Supplemental Figure 2A, in which different ancestries had different levels of private germline variants within a fixed filtering regime, our models improved in their performance metrics only in the context of tumor-only data when adding ancestry.

### Supplemental Data Description

The panel inputs in Data_S1 contain all variants which fell within the coordinates of the respective panels and coverage WIGs and CDS and divided by the covered panel CDS size in Mb. As a result these counts include synonymous mutations as well as hotspots. Because germline data is potentially sensitive information we anonymized the sample names. Samples which had less than 400 germline counts in the 4 Mb region prior to filtering were excluded from the germline inputs.

### Supplemental Figures

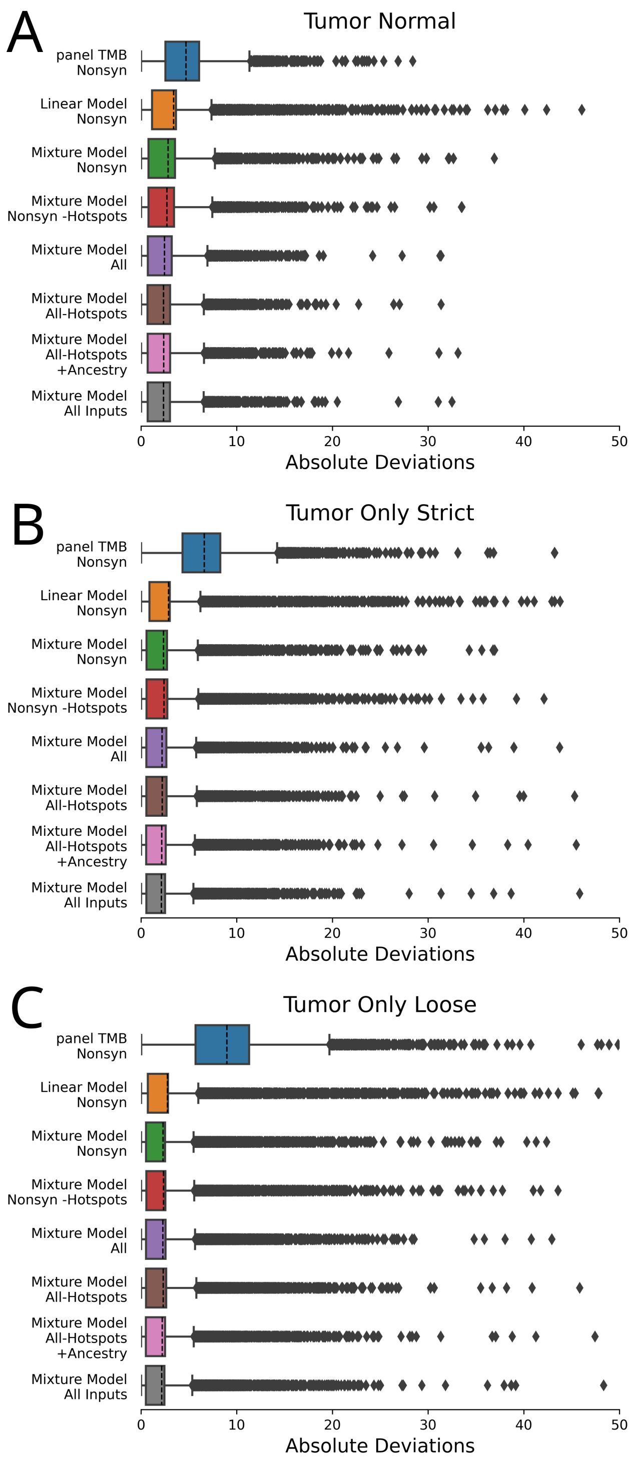


**Supplemental Figure 1. Box plots of the data represented in Table 1.** Mean of the deviations represented as a dashed line.

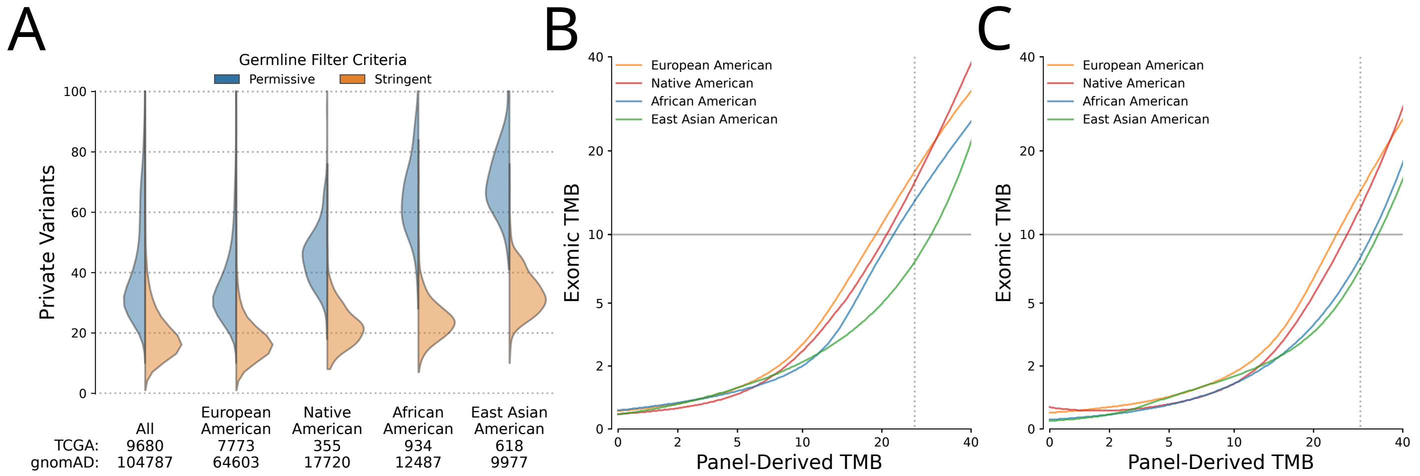


**Supplemental Figure 2. Effect of ancestry on germline filtering and model fits.** (A) Numbers of germline variants which passed the different filtering criteria for different ancestries. (B) Model fits for models trained on the different cohorts for stringent tumor-only data. (C) Model fits for models trained on the different cohorts for permissive tumor-only data.

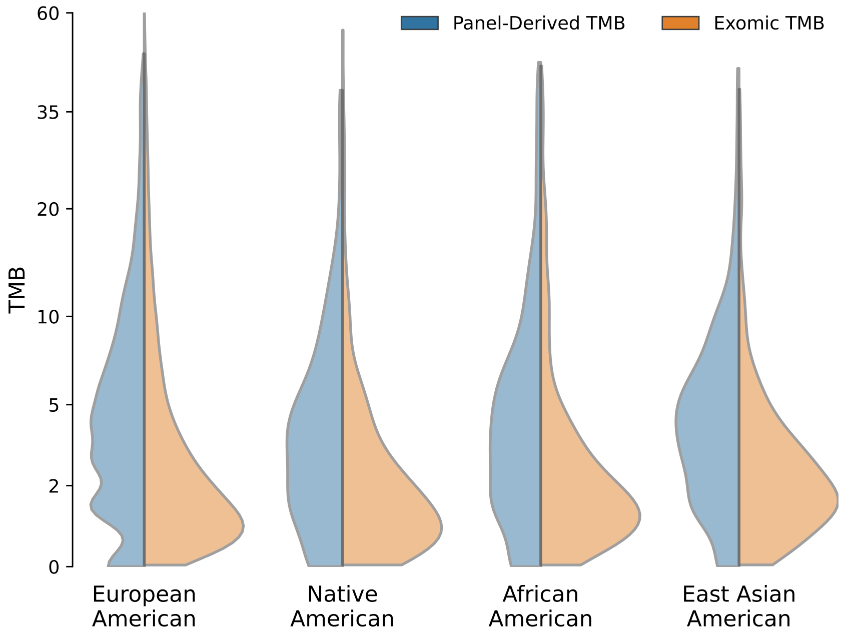


**Supplemental Figure 3. TMB distributions of different ancestries.**

### Supplemental Tables

| | | Tumor Normal | | Tumor Only | |
|---|---|---|---|---|---|
| | | Exomic TMB $> t$ | n | Exomic TMB $> t$ | n |
| Panel TMB | $\hat{y} > t$ | 53.5% | 1486 | 30.6% | 2758 |
| | $\hat{y} \leq t$ | 1.1% | 8201 | 0.5% | 6838 |
| Linear Model | $\hat{y} > t$ | 94.3% | 507 | 93.1% | 379 |
| | $\hat{y} \leq t$ | 4.4% | 9180 | 5.7% | 9217 |
| | $>$ 95% certain, $\hat{y} > t$ | 100.0% | 102 | 0.0% | 0 |
| | $>$ 95% certain, $\hat{y} < t$ | 0.5% | 7733 | 0.6% | 7075 |
| | $<$ 95% certain, $\hat{y} \neq t$ | 40.1% | 1852 | 33.3% | 2521 |
| Mixture Model | $\hat{y} > t$ | 87.7% | 661 | 81.5% | 653 |
| | $\hat{y} \leq t$ | 3.3% | 9026 | 3.9% | 8943 |
| | $>$ 95% certain, $\hat{y} > t$ | 99.6% | 247 | 94.7% | 171 |
| | $>$ 95% certain, $\hat{y} < t$ | 0.5% | 7716 | 0.8% | 7415 |
| | $<$ 95% certain, $\hat{y} \neq t$ | 34.8% | 1724 | 32.5% | 2010 |

**Supplemental Table 1. Detection rates of somatic, exomic TMB $>$ 10 (t) based on stratification by panel-derived TMB measures.** The detection rate of the observed true somatic, exomic TMB being greater than the threshold (t) of 10 based on the various panel derived measures as designated. The linear model and mixture model values are derived from the data in Figure 1 of the manuscript. $\hat{y}$: model prediction

| Data | Ancestry | With Ancestry | MAE | Spearman's rho |
|---|---|---|---|---|
| Tumor Normal (matched germline substracted) | European American | - | 2.45 | 0.81 |
| | | + | 2.40 | 0.82 |
| | Native American | - | 1.73 | 0.83 |
| | | + | 1.84 | 0.79 |
| | African American | - | 2.14 | 0.78 |
| | | + | 2.17 | 0.78 |
| | East Asian American | - | 1.87 | 0.68 |
| | | + | 1.95 | 0.67 |
| Tumor Only (stringent germline filtering) | European American | - | 2.45 | 0.67 |
| | | + | 2.27 | 0.67 |
| | Native American | - | 1.69 | 0.62 |
| | | + | 1.68 | 0.61 |
| | African American | - | 1.88 | 0.59 |
| | | + | 1.88 | 0.59 |
| | East Asian American | - | 1.62 | 0.56 |
| | | + | 1.57 | 0.55 |
| Tumor Only (permissive germline filtering) | European American | - | 2.50 | 0.58 |
| | | + | 2.30 | 0.59 |
| | Native American | - | 1.73 | 0.52 |
| | | + | 1.71 | 0.51 |
| | African American | - | 1.82 | 0.56 |
| | | + | 1.81 | 0.56 |
| | East Asian American | - | 1.66 | 0.49 |
| | | + | 1.54 | 0.50 |

**Supplemental Table 2. TMB regression metrics for different ancestries** All metrics are from cross validation with counts of all mutations, hotspots, and nonsynonymous mutations as the input. MAE: mean absolute error. MAE and Spearman rank-order correlation are calculated for samples with panel TMB greater than or equal to 5.
